# Meiotic MCM proteins promote and inhibit crossovers during meiotic recombination

**DOI:** 10.1101/467134

**Authors:** Michaelyn Hartmann, Kathryn P. Kohl, Jeff Sekelsky, Talia Hatkevich

## Abstract

Crossover formation as a result of meiotic recombination is vital for proper segregation of homologous chromosomes at the end of meiosis I. In many organisms, crossovers are generated through two crossover pathways: Class I and Class II. To ensure accurate crossover formation, meiosis-specific protein complexes regulate the degree in which each pathway is used. One such complex is the mei-MCM complex, which contains MCM (mini-chromosome maintenance) and MCM-like proteins REC (ortholog of Mcm8), MEI-217, and MEI-218, collectively called the mei-MCM complex. The mei-MCM complex genetically promotes Class I crossovers and inhibits Class II crossovers in *Drosophila*, but it is unclear how individual mei-MCM proteins contribute to crossover regulation. In this study, we perform genetic analyses to understand how specific regions and motifs of mei-MCM proteins contribute to Class I and II crossover formation and distribution. Our analyses show that the long, disordered N-terminus of MEI-218 is dispensable for crossover formation, and that mutations that disrupt REC’s Walker A and B motifs differentially affect Class I and Class II crossover formation. In Rec Walker A mutants, Class I crossovers exhibit no change, but Class II crossovers are increased. However, in *rec* Walker B mutants, Class I crossovers are severely impaired, and Class II crossovers are increased. These results suggest that REC may form multiple complexes that exhibit differential REC-dependent ATP binding and hydrolyzing requirements. These results provide genetic insight into the mechanisms through which mei-MCM proteins promote Class I crossovers and inhibit Class II crossovers.

## Introduction

To reestablish the diploid genome upon sexual fertilization, the genome of progenitor germ cells must be successfully reduced by half through meiosis. Accurate reduction of the genome at the end of meiosis I requires crossover formation between homologous chromosomes during meiotic recombination. Meiotic recombination is initiated by the formation of multiple double-strand breaks (DSBs); the majority of meiotic DSBs are repaired as noncrossovers, while a selected subset are repaired as crossovers between homologs (reviewed in Lake and Hawley 2012).

Two distinct types of meiotic crossovers have been described: Class I and Class II. First defined in budding yeast (de Los Santos *et al.* 2003), Class I and Class II crossovers exist in most sexually reproducing organisms, but the relative proportions of each crossover type vary among organisms (Hollingsworth and Brill 2004). In *Drosophila*, most – if not all – crossovers are generated through the Class I pathway (Hatkevich *et al.* 2017), as shown through their dependence on the putative catalytic unit of the Class I meiotic resolvase MEI-9 (Sekelsky *et al.* 1995; Yildiz *et al.* 2002) and their display of crossover interference (Hatkevich *et al.* 2017). Most crossovers in *Drosophila* are also dependent upon a group of MCM- or MCM-like proteins, called the mei-MCM complex (Baker and Carpenter 1972; Grell 1978; Liu *et al.* 2000; Kohl *et al.* 2012).

The mei-MCM complex consists of REC (the *Drosophila* ortholog of MCM8), MEI-217, and MEI-218. REC appears to be a bona fide MCM protein, based on conservation of both the N-terminal MCM domain and the C-terminal AAA+ ATPase domain, which includes Walker A and B boxes that bind and hydrolyze ATP (Figure 1A). In contrast, MEI-217 and MEI-218 are highly divergent MCM-like proteins, and together resemble one full MCM protein. MEI-217 is structurally similar to the MCM N-terminal domain, though this similarity is not detected in BLAST or conserved domain searches (Kohl *et al.* 2012). The carboxy-terminus of MEI-218 has a domain related to the AAA+ ATPase domain, but key residues are not conserved, including the Walker A and B motifs that are critical for binding and hydrolyzing ATP, respectively (Iyer *et al.* 2004) (Figure 1B). Because key residues in the Walker A and B motifs are not conversed, MEI-218 may not exhibit ATPase activity or it may exhibit partial function. In addition, MEI-218 has a long N-terminal extension that is poorly conserved and predicted to be disordered. The function of this region is unknown, but gene swap studies suggest that it may contribute to differences in the recombination landscape among *Drosophila* species (Brand *et al.* 2018). For further analysis and details regarding the evolution of the mei-MCM complex, see Supplemental Figures S1-S3.

**Figure 1.**
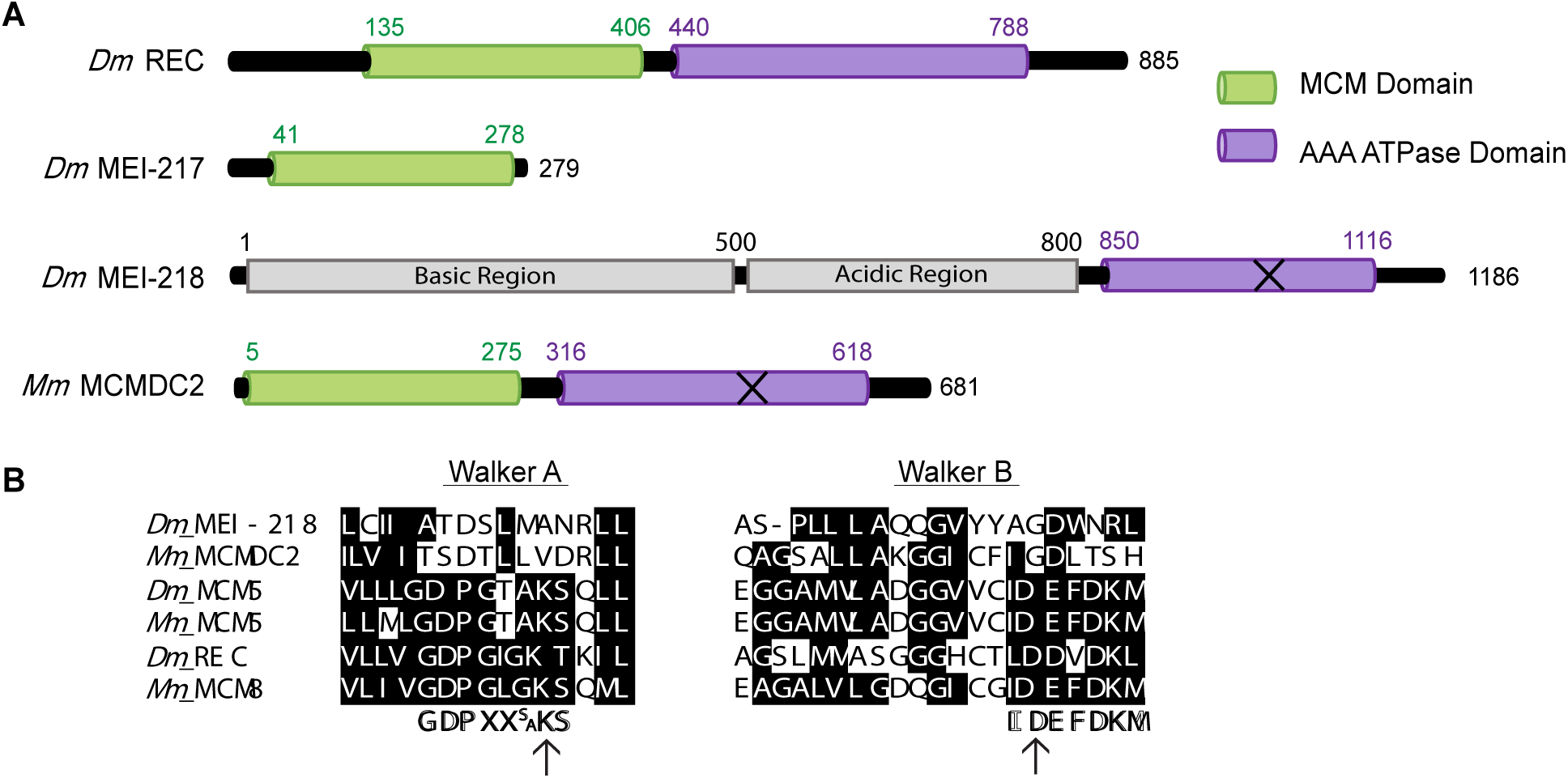
MCM protein structure and alignments. (A) Structural domains of *Drosophila melanogaster* REC, MEI-217, MEI-218 and *Mus musculus* MCMDC2. Structural domains identified using PHYRE 2 (Kohl *et al.* 2012). “MCM domain” corresponds to protein data bank ID #c2vl6C and the AAA ATPase domains identified correspond to protein data bank ID #d1g8pa. The X on *Dm* MEI-218 and *Mm* MCMDC2 represents predicted inactive AAA ATPase domains. (B) Consensus sequence for Walker A motif (Walker *et al.* 1982), and consensus sequence for Walker B motif (Forsburg 2004). Identical or conserved amino acids are denoted with black background. Arrows denote the conserved catalytic residues.

While most crossovers are generated through the Class I pathway in wild-type *Drosophila* and are mei-MCM dependent, mutants that lack the Bloom syndrome helicase (Blm) generate only Class II crossovers based on their independence of MEI-9 and lack of patterning (*e.g.*, interference) that is associated with Class I crossovers (Hatkevich *et al.* 2017). Blm is an ATP-dependent 3’-5’ helicase that exhibits vital anti-crossover functions in both meiotic and somatic DSB repair (reviewed in Hatkevich and Sekelsky 2017). Interestingly, mutations in *mei-MCM* and *Blm* genes genetically interact. In *Blm* mutants, crossovers are reduced by 30% but in a *Blm rec* double mutant, crossovers are significantly increased compared to wild-type (Kohl *et al.* 2012). This suggests that the mei-MCMs may function to inhibit crossovers within the Class II pathway, in addition to their role promoting crossovers in the Class I pathway.

While the mei-MCMs function as a complex, little is known about how individual mei-MCMs contribute to Class I and II crossover regulation. Here, we investigate specific features of MEI-218 and REC to understand better how these proteins contribute to meiotic recombination. We find that the N-terminus of MEI-218 is dispensable for crossover formation and general crossover distribution. By mutating key residues in REC’s Walker A and B motifs (*rec*^*KA*^ and *rec*^*DA*^, respectively), we found that *rec*^*KA*^ mutants exhibit no Class I crossover defect, while Class II crossovers are significantly increased. Surprisingly, *rec*^*DA*^ mutants exhibit a severe decrease in Class I crossovers and a significant increase in Class II crossovers. Our results suggest that the mei-MCMs function in multiple roles and may complex in a variety of configurations to properly regulate crossover formation.

## Materials and Methods

### Drosophila stocks

Flies were maintained on standard medium at 25°C. Some mutant alleles have been previously described, including *mei-9*^*a*^ (Baker and Carpenter 1972; Yildiz *et al.* 2004), *mei-218*^*1*^ and *mei-218*^*6*^ (Baker and Carpenter 1972; Mckim *et al.* 1996), *Blm*^*N1*^ and *Blm*^*D2*^ (Mcvey *et al.* 2007), *rec*^*1*^ and *rec*^*2*^ (Grell 1978; Matsubayashi and Yamamoto 2003; Blanton *et al.* 2005). The maternal-effect lethality in *Blm*^*N1*^*/Blm*^*D2*^ mutants was overcome by the *UAS::GAL4* rescue system previously described (Kohl *et al.* 2012).

### Generating mei-218 transgenic alleles

The transgenes for *mei-218Δ*^*N*^ and *mei-218*^*FL*^ were constructed by cloning cDNA for *mei-218* into *P*{*attBUASpW*} (AddGene). Full-length *mei-218* included codons 1-1186; the *mei-218Δ*^*N*^ transgene included codons 527-1186. Transgenics were made by integrating into a phiC31 landing site in 2A on the *X* chromosome.

### *Generating* rec^KA^ *and* rec^DA^ *mutants*

Annealed oligonucleotides were inserted into *Bbs*I-digestedpU6-BbsI-chiRNA plasmid(Addgene). *rec*^*KA*^: CTTCGCCGAGAAGGGATAGTAAAC; *rec*^*DA*^: CTTCGTTGCAGTGCCTACAATCAG. Resulting plasmids were co-injected with repair template plasmid, consisting of synthesized gBlocks (IDT DNA) cloned into pBlueScript plasmid (sequences available on request). Injected larvae were raised to adulthood, and their male progeny were crossed to *TM3*/*TM6B* females (Bloomington Stock Center) to generate stocks, after which DNA was extracted for screening through PCR and restriction digest.

### Nondisjunction assay

*X*-chromosome nondisjunction (NDJ) was assayed by mating virgin females to *y cv v f* / *T(1:Y)B*^*S*^ males. Each cross was set up as a single experiment with 20-50 separate vials. The progeny of each vial were counted separately. Viable nondisjunction progeny are *XXY* females with Bar eyes and *XO* males with Bar^+^ eyes and the phenotypes from *y cv v f* chromosome. Total (adjusted) represents the total with inviable exceptional progeny accounted for (*XXX* and *YO*). NDJ rates and statistical comparisons were done as in Zeng et al. 2010.

### Crossover distribution assay

Crossover distribution on chromosome *2*L was scored by crossing virgin *net dpp*^*d-ho*^ *dp b pr cn / +* female flies with mutant background of interest to *net dpp*^*d-ho*^ *dp b pr cn* homozygous males. Each cross was set up as a single experiment with at least 25 separate vials scored. The first set of vials was flipped after three days of mating into vials of a new batch, although these were counted as one experiment. Batch effects for recombination assays have not been observed in repeated studies for multiple genotypes used in this study (Figure S4). These include wild-type (unpublished data), *Blm* (unpublished data), *rec* (Blanton *et al.* 2005; Kohl *et al.* 2012), *mei-9* (Sekelsky *et al.* 1995), and *mei-9; rec* (Blanton *et al.* 2005). All progeny were scored for parental and recombinant phenotypes. Crossover numbers in flies are shown as cM where cM = (number of crossovers / total number of flies) * 100. Chi-squared tests with Bonferroni correction were performed for each interval. For total cM, Fisher’s Exact Test was used to compare total crossovers to total number of flies. Crossover distribution is represented as cM/Mb where Mb is length of the interval without transposable elements (TEs) because crossovers rarely occur within TEs (Miller *et al.* 2016).

### Protein structure and alignment

Structural domains of proteins were determined by using PHYRE 2. All of the MCM regions identified correspond to the protein data bank ID #c2vl6C and the AAA ATPase domains identified correspond to protein data bank ID #d1g8pa. Alignment of the Walker A and Walker B motifs (Kohl *et al.* 2012) was done using MEGA 5 and aligned with the ClustalW program. Identical and conserved residues are shaded based on groups of amino acids with similar chemical properties.

### Data availability

All data necessary for confirming the conclusions in this paper are included in this article and in supplemental figures and tables. *Drosophila* stocks and plasmids described in this study are available upon request. We have uploaded Supplemental Material to Figshare. Figure S1 illustrates distribution of Msh4, Msh5, Mcm8, Mcm9, MEI-217, and MEI-218 in Diptera. Figure S2 illustrates the structure of MEI-217 and MEI-218 in Diptera. Figure S3 shows sequence alignment of MEI-218. Figure S4 compares crossover frequencies in different batches of the same genotype. Figure S5 details the cross scheme of *mei-218* transgene experiments. Table S1 includes analysis of genetic interval differences between *WT* and *mei-218*^*FL*^. Table S1 includes analysis of genetic interval differences between *mei-218*^*FL*^ and *mei-218*^*ΔN*^. Table S2 includes complete data set for calculating non-disjunction of *WT, rec*^*-*^*/rec*^*+*^, and *rec*^*DA*^*/+.* Table S3 includes all data sets for meiotic crossovers for all genotypes discussed.

## Results and Discussion

### The N-terminus of MEI-218 is dispensable for crossover formation

MCMDC2 is a distantly-related member of the MCM family of proteins that is unique in that the ATPase domain is predicted to be incapable of binding or hydrolyzing ATP. Orthologs in Dipteran insects are further distinguished by having the N-terminal and ATPase-like domains encoded in separate open reading frames. The two polypeptides, MEI-217 and MEI-218 interact physically, at least in *Drosophila melanogaster*, presumably reconstituting a single MCM-like protein. MEI-218 is also distinguished by possessing an N-terminal extension of variable length in different species. *Drosophila melanogaster* MEI-218 can be divided into three distinct regions (Figure 1A): an N-terminal tail (residues 1-500), a central acidic region (residues 500-800) and the C-terminal ATPase-related region (residues 850-1116) (Kohl *et al.* 2012; Brand *et al.* 2018). The N-terminal and middle regions are predicted to be disordered (Kohl *et al.* 2012) and are poorly conserved (Figure S3). Results obtained through gene swap experiments suggest that the N-terminal tail and central region regulate crossover number and distribution within *Drosophila* species (Brand *et al.* 2018).

To genetically examine the function of the N-terminus of MEI-218, we compared functions of a transgene that expresses a truncated form of MEI-218 that lacks the N-terminal 526 amino acids (*mei-218Δ*^*N*^) to a matched full-length transgene (*mei-218*^*FL*^) (Figure 2A). Due to the relatively high conservation among *Drosophila* species, the middle region of mei-218 was retained for this experiment (Figure S3). Using the *UAS*/*GAL4* system (Duffy 2002), we expressed both constructs in *mei-218* null mutants using the germline-specific *nanos* promoter and measured crossovers along five adjacent intervals that span most of *2*L and part of *2R* (Figure S4; for simplicity, we refer to this chromosomal region as *2*L.)

**Figure 2.**
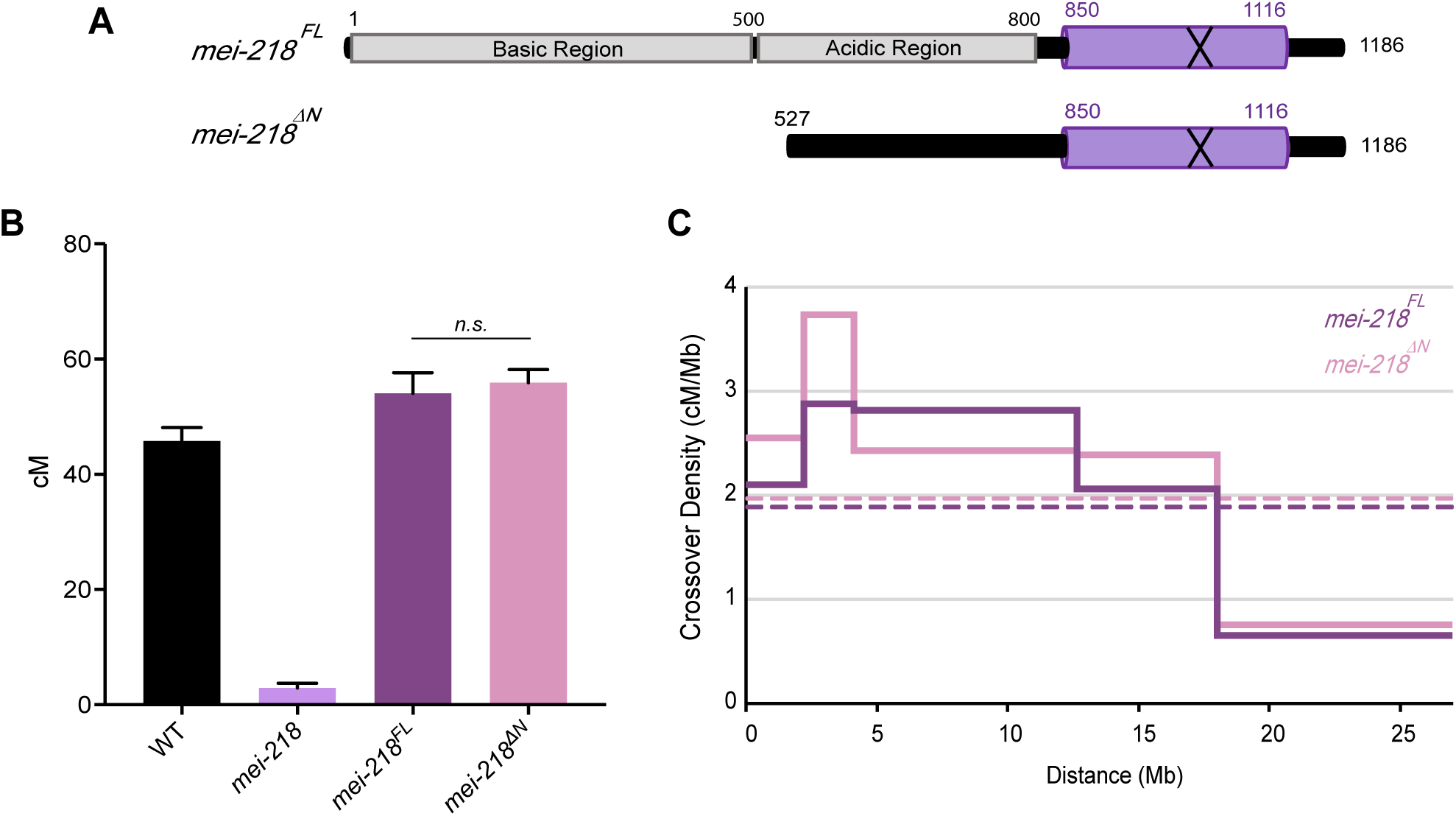
The role of MEI-218 N-terminus in crossover formation and distribution. (A) Schematic of transgenes for full length *mei-218* and N-terminal deleted *mei-218*, in which the first 526 amino acids are absent. (B) Map units of WT (Hatkevich *et al.* 2017), *mei-218* (Kohl *et al.* 2012), *mei-218*^*FL*^ and *mei-218*^ΔN^. Map units represented as centimorgans (cM). Error bars indicate 95% confidence intervals. *n.s. =* not significant (*p* = 0.61). (C) Crossover distribution (solid lines) of *mei-218*^*FL*^ and *mei-218*^ΔN^ represented as cM/Mb. Mb is measured distance of defined interval, excluding centromere, pericentromeric heterochromatin and transposable elements. Dotted lines represent mean crossover density across *2*L. Figure S5 details the cross scheme of *mei-218* transgene experiments. Refer to tables S1and S3 for complete data sets.

In wild-type females, the genetic length of *2*L is 45.8 cM (Hatkevich *et al.* 2017) (Figure 2B), whereas *mei-218* mutants exhibit a severe decrease in crossovers, with genetic length of 2.92 cM (Kohl *et al.* 2012). Expression of *mei-218*^*FL*^ in *mei-218* mutants (*mei-218*^*FL*^) fully rescues the crossover defect, exhibiting a genetic length of 54.1 cM. Unexpectedly, expression of *mei-218Δ*^*N*^ in *mei-218* mutants (*mei-218Δ*^*N*^) restored crossing over to the same level as in *mei-218; mei-218*^*FL*^ (55.9 cM; *n.s. p* = 0.61).

Brand *et al.* (2018) expressed *Drosophila mauritiana* MEI-217 and MEI-218 in *Drosophila melanogaster* and found that crossovers were increased in proximal and distal regions, resulting in an overall change in crossover distribution. We examined crossover distribution in *mei-218; mei-218*^*FL*^ and *mei-218; *mei-218Δ*^*N*^* (Figure 2C). Overall, distributions are similar, with both genotypes exhibiting a strong inhibition of crossovers near the centromere (referred to as the centromere effect; Beadle 1932) and the majority of the crossovers placed in the medial-distal regions (Figure 2C).

We conclude that the N-terminal tail of MEI-218 is dispensable for both crossover formation and overall distribution on chromosome *2*L. This conclusion is supported by the observation that of 16 sequenced mutations in *Drosophila melanogaster mei-218*, 14 are nonsense or frameshift, and the only two missense mutations alter residues in the C-terminus (amino acids 845 and 1107) (Collins *et al.* 2012).

The reasons why the MCM domains have been separated into MEI-217 and MEI-218 polypeptides and why MEI-218 has an N-terminal extension are unknown, but this structure has been maintained for more than 250 million years of Dipteran evolution (Supplemental Figures S2). Interestingly, MEI-218 is expressed moderately highly in testes (Thurmond *et al.* 2018) even though males do not experience meiotic recombination. The predominant or exclusive transcript in males does not encode MEI-217 (Thurmond *et al.* 2018), the seemingly obligate partner for MEI-218 the female meiotic recombination. Males that lack *mei-218* are viable, fertile, and do not exhibit elevated nondisjunction (Baker and Carpenter 1972; Mckim *et al.* 1996). For these reasons, we speculate that an unknown function of MEI-218 (independent of MEI-217) in the male germline explains why its overall structure has been evolutionarily maintained.

### REC ATPase motifs are required for crossover formation

Of the three known mei-MCM subunits, only REC harbors well-conserved Walker A and B motifs, suggesting that REC has ATP binding and hydrolysis activity (Kohl *et al.* 2012). It is unknown whether the mei-MCM complex utilizes REC’s putative ATPase activity for its function *in vivo*. To test this, we used CRIPSR/Cas9 to introduce into *rec* mutations predicted to disrupt Walker A and B motif functions (Figure 3A). The Walker A mutation (*rec*^*KA*^) results in substitution of a conserved lysine residue with alanine; this mutation in other AAA+ ATPases, including replicative MCMs, prevents binding of ATP (Bell and Botchan 2013). The Walker B mutation (*rec*^*DA*^) results in substitution of a conserved aspartic acid with alanine; in MCMs and other AAA+ ATPases, this mutation destroys the ability to coordinate Mg^++^ for ATP hydrolysis (Bochman *et al.* 2008).

**Figure 3.**
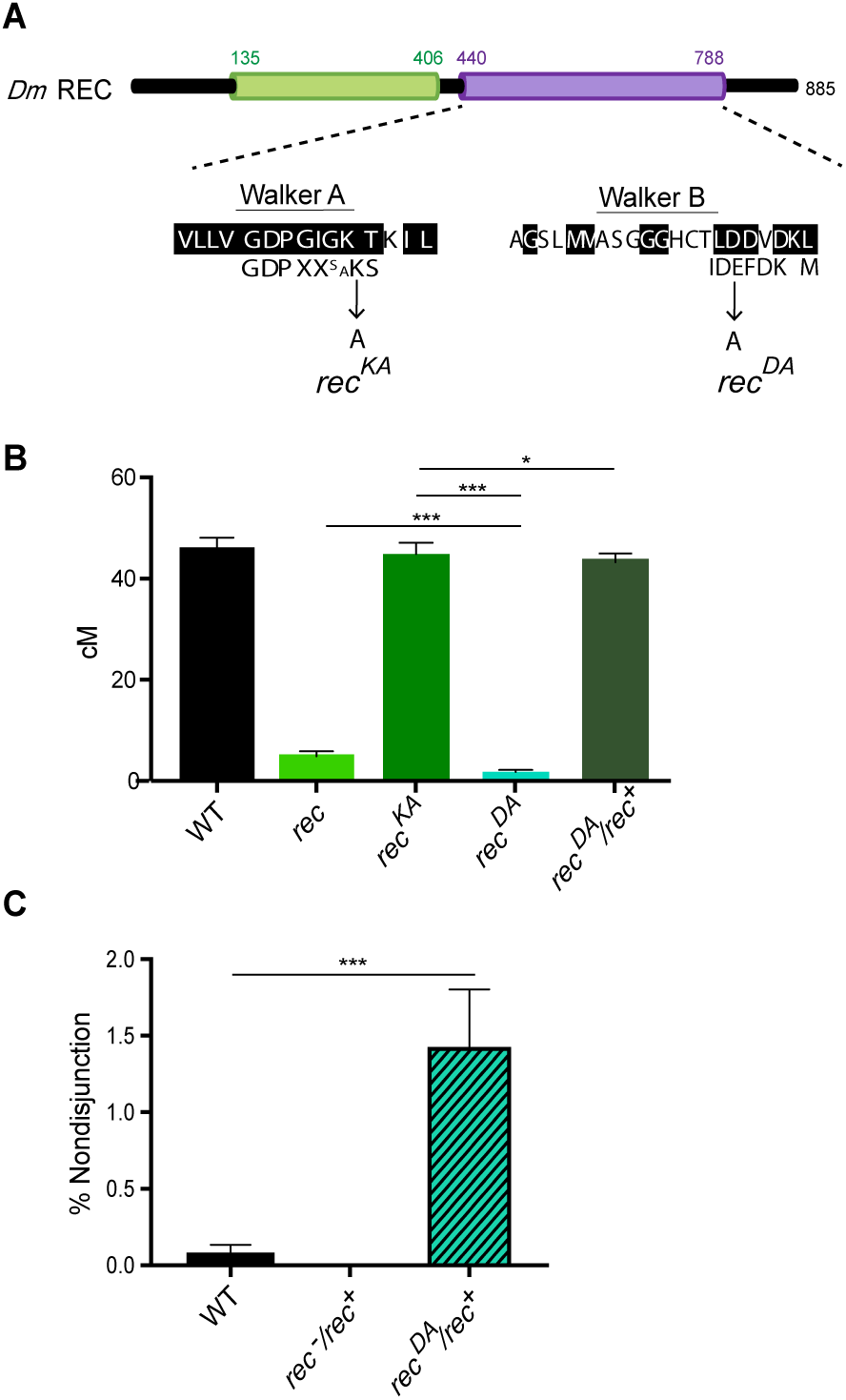
REC ATPase binding and hydrolysis requirements for crossover formation. (A) Schematic representation of the mutated residues in *rec*^*KA*^ and *rec*^*DA*^. (B) Map units of *WT* (Hatkevich *et al.* 2017), *rec*^*1*^*/rec*^*2*^, *rec*^*KA*^, and *rec*^*DA*^, *rec*^*DA*^*/rec*^*+*^. Map units represented as centimorgans (cM). Error bars show 95% confidence intervals. (C) Percent nondisjunction of *WT, rec*^*1*^*/rec*^*+*^, and *rec*^*DA*^*/rec*^*+*^. (D) Model of possible complex depicting the functional Walker B motif of REC protein interacting with a Walker A motif on a potential partner. * *p* < 0.05; *\*\*\*p <* 0.0001. Refer to tables S2 and S3 for complete data sets.

We assayed crossover frequency along *2*L in *rec*^*KA*^ and *rec*^*DA*^ mutants (Figure 3B). Surprisingly *rec*^*KA*^ ATP binding mutants exhibit a genetic length of 44.9 cM, which is not significantly different from wild-type (*p* = 0.4016), suggesting that ATP binding by REC is not required for crossover formation. Conversely, there is a severe reduction in crossovers in *rec*^*DA*^ mutants, with a genetic length of 1.6 cM (*p* < 0.0001), suggesting that REC’s ability to hydrolyze ATP is required for crossover formation.

Because the genetic length of *rec*^*DA*^ is significantly lower than *rec* null mutants (Figure 3B, *p* < 0.0001), we hypothesized that *rec*^*DA*^ is an antimorphic mutation. To test this, we examined crossover levels and *X* chromosome nondisjunction (NDJ) in *rec*^*DA*^/*rec*^*+*^ (Figure 3B and 3C, respectively). The genetic length of *2*L in *rec*^*DA/+*^ is slightly lower than wild-type, but not significantly different (43.9 cM and 45.8 cM, respectively; *p* = 0.35). For *X*-NDJ, both wild-type and *rec*^*-*^*/rec*^*+*^ mutants exhibit rates below 0.5%, while *rec*^*DA*^*/rec*^*+*^ mutants exhibit a significant increase to 1.4% NDJ (*p* < 0.0001). These data support the conclusion that *rec*^*DA*^ is weakly antimorphic and suggest that *rec*^*DA*^ results in an inactive mei-MCM complex that is antagonistic to the wild-type complex. In light of these interpretations, we propose that the mei-MCM complex binds to recombination sites independent of REC binding to ATP, and that REC-dependent ATP hydrolysis is required for the removal of the mei-MCM complex from these sites.

The phenotypes of *rec*^*KA*^ and *rec*^*DA*^ mutants suggest that REC’s ability to hydrolyze ATP is required for crossover formation, whereas its ATP binding capability is dispensable. The disparate requirements for REC’s ATP binding and hydrolysis are similar to those of other ATPase-dependent complexes. Rad51 paralogs, which form multi-protein complexes and contain Walker A and B motifs, are proposed to exhibit ATPase activity in *trans* between adjacent subunits, each of which contributes a Walker A or Walker B motif to the active site (Wu *et al.* 2004; Wu *et al.* 2005; Wiese *et al.* 2006). Because neither MEI-217 nor MEI-218 possess an ATPase domain that harbors conserved key enzymatic residues (Figure 1B) (Kohl *et al.* 2012), we propose that ATPase activity of the mei-MCM complex requires REC for ATP hydrolysis and an unknown mei-MCM protein for ATP binding. Alternatively, because REC is highly diverged, its Walker A and B motifs may function non-canonically. Biochemical studies are needed to test these hypotheses, but these may require identification of the putative missing subunit.

### REC-dependent ATP hydrolysis is required for MEI-9-dependent crossovers

To gain insight into the crossover pathways that are used in *rec*^*KA*^ and *rec*^*DA*^ mutants, we examined whether these crossovers require the Class I endonuclease/resolvase. In *Drosophila*, the catalytic subunit of the putative Class I meiosis-specific endonuclease is MEI-9 (Sekelsky *et al.* 1995; Yildiz *et al.* 2002; Hatkevich *et al.* 2017). The *2*L genetic length within a *mei-9* mutant is 2.75 cM (Figure 4), demonstrating that at least 90% of crossovers are dependent upon MEI-9. However, the genetic length in *mei-9*; *rec* mutants is not significantly different than that of *rec* null single mutants (4.11 cM *vs* 4.66 cM, *p* = 0.64) suggesting that in the absence of REC, the resulting crossovers are likely independent of MEI-9. Similarly, it has been shown previously that *mei-218 mei-9* double mutants do not have reduced crossovers compared to *mei-218* single mutants (Sekelsky *et al.* 1995), indicating that crossovers generated in the absence of the mei-MCM complex are MEI-9-independent.

**Figure 4.**
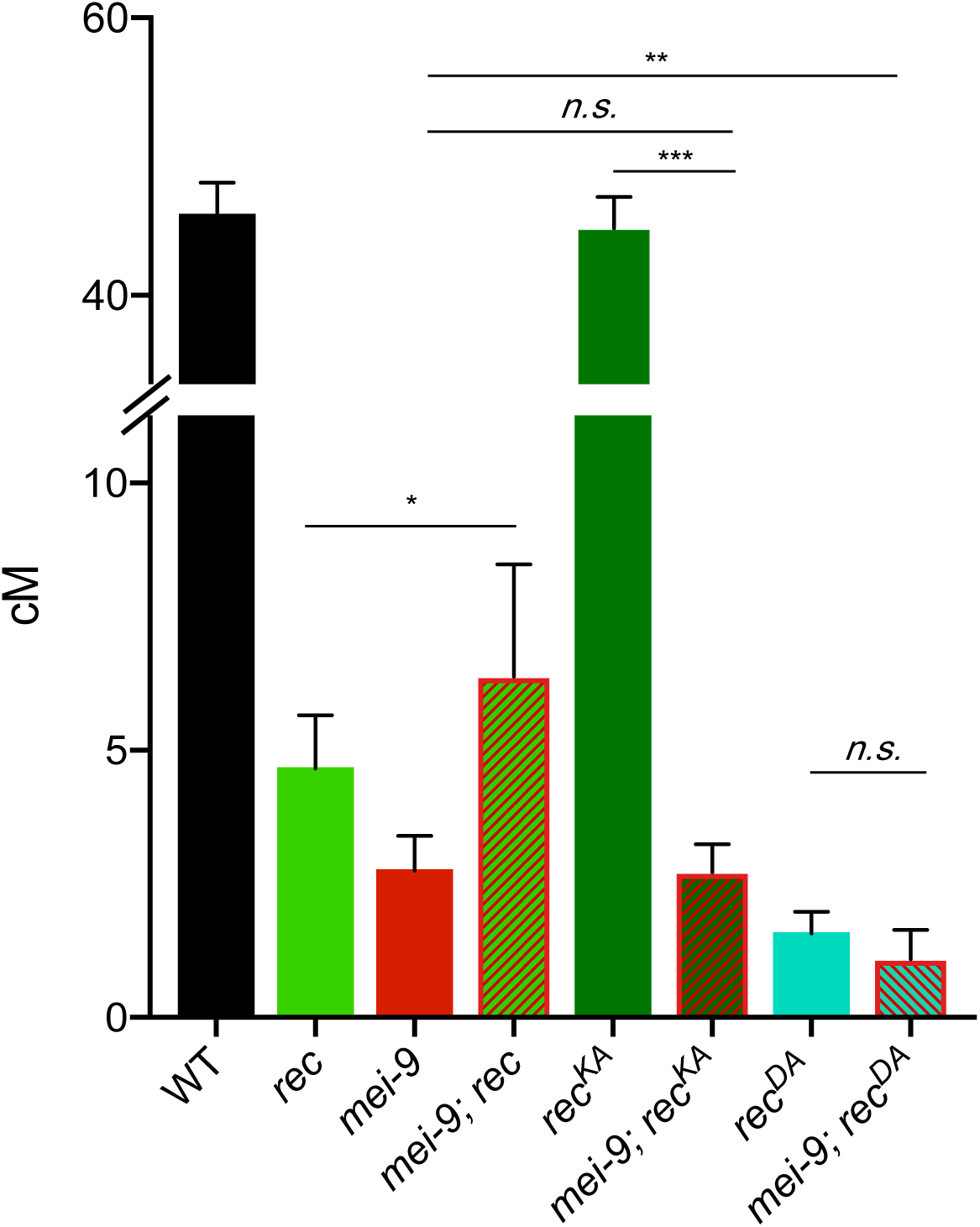
MEI-9-dependent crossovers in *rec*^*KA*^ and *rec*^*DA*^ mutants. Map units of *WT* (Hatkevich *et al.* 2017), *rec, mei-9, mei-9;rec, rec*^*KA*^, *mei-9;rec*^*KA*^, *rec*^*DA*^, and *mei-9;rec*^*DA*^. Map units represented as centimorgans (cM). Error bars show 95% confidence intervals. * *p* < 0.05 ** *p* < 0.001 \*\*\**p* < 0.0001; (*mei-9* vs *mei-9; rec*^*KA*^ *p* = 0.94) (*rec*^*DA*^ vs *mei-9; rec*^*DA*^ *p* = 0.23). Refer to Table S3 for complete data set.

Because *rec*^*KA*^ mutants exhibit the same distribution and number of crossovers as wild-type (Figure 3B), we hypothesized that *rec*^*KA*^ crossovers are dependent on MEI-9. To test this, we examined genetic length across *2*L in *mei-9; rec*^*KA*^ double mutants (Figure 4). Mutants for *mei-9; rec*^*KA*^ exhibit a genetic length of 2.72 cM, which is significantly decreased compared to the *rec*^*KA*^ single mutant (*p* < 0.0001), but not significantly different from *mei-9* single mutants (*p* = 0.94), showing that crossovers in *rec*^*KA*^ are indeed dependent upon MEI-9 nuclease. In contrast, we predicted that crossovers in *rec*^*DA*^ will be independent of MEI-9, similar to crossovers generated in *rec* null mutants. We observe that *mei-9; rec*^*DA*^ double mutants exhibit a genetic length of 1.1 cM, which is significantly lower than that of *mei-9* single mutants (*p* < 0.001). Importantly, crossing over in the *mei-9; rec*^*DA*^ double mutant is not significantly different than in *rec*^*DA*^ single mutants (*p* = 0.23), demonstrating that crossovers in *rec*^*DA*^ are independent of MEI-9 (Figure 4).

From these data we conclude that the crossovers in *rec*^*KA*^ mutants arise through the normal, MEI-9-dependent pathway, whereas mitotic nucleases generate the residual crossovers in *rec*^*DA*^ mutants. These data show that REC^KA^ functions normally in the Class I pathway, but this pathway is nonfunctional in *rec* null and *rec*^*DA*^ mutants. We suggest that the REC’s ability to hydrolyze, but not bind, ATP is required for the formation of Class I crossovers.

### REC ATPase motifs are required to prevent Class II crossovers

In wild-type *Drosophila*, most or all crossovers are generated through the Class I pathway (Hatkevich *et al.* 2017), and these crossovers are dependent upon the mei-MCM complex (Kohl *et al.* 2012). However, in *Blm* mutants, crossovers are generated exclusively through the Class II pathway (Hatkevich *et al.* 2017). In *Drosophila Blm* mutants, meiotic crossovers are decreased by 30%, suggesting that the Class II pathway is less efficient at generating crossovers than the Class I pathway, even though what may be the primary anti-crossover protein, Blm helicase, is absent. It has previously been shown that loss of Blm suppresses the high nondisjunction of *mei-218* and *rec* mutants (Kohl *et al.* 2012). However, in *Blm rec* double mutants, crossovers are increased significantly compared to *Blm* single mutants (Kohl *et al.* 2012), suggesting that REC and/or the mei-MCM complex has an anti-crossover role in *Blm* mutants, and therefore in the Class II crossover pathway.

To further understand the role of REC in the Class II pathway, we investigated whether REC’s predicted ATP binding or hydrolysis function is required for its Class II anti-crossover function. To do this, we measured the crossovers across *2*L in *rec*^*KA*^ and *rec*^*DA*^ in the background of *Blm* mutants. If REC ATP binding or hydrolysis is required for an anti-crossover role in Class II, then the genetic length of *Blm rec*^*KA*^ or *Blm rec*^*DA*^ double mutants will be similar to that of *Blm rec* double mutants. Conversely, if REC ATP binding or hydrolysis is not required, then double mutants will exhibit genetic lengths similar to that of *Blm* single mutants.

Interestingly, *Blm rec*^*KA*^ mutants exhibit a genetic length of 43.3 cM, which is not significantly different than *Blm rec* mutants (*p* = 0.10) but significantly higher than *Blm* single mutants (*p* < 0.0001; Figure 5A). Similarly, *Blm rec*^*DA*^ double mutants have a recombination rate of 53.4 cM, which not significantly different from *Blm rec* double mutants (*p* = 0.52), but significantly higher than *Blm* single mutants (*p* < 0.0001). These results suggest that REC’s predicted abilities to bind and hydrolyze ATP are both required for the inhibition of crossovers at REC-associated Class II recombination sites. Therefore, it appears that REC forms different complexes within the Class II pathway and Class I pathway. It is unknown whether this Class II REC-associated complex requires the other mei-MCM proteins, and additional genetic studies will be valuable to discern this.

**Figure 5.**
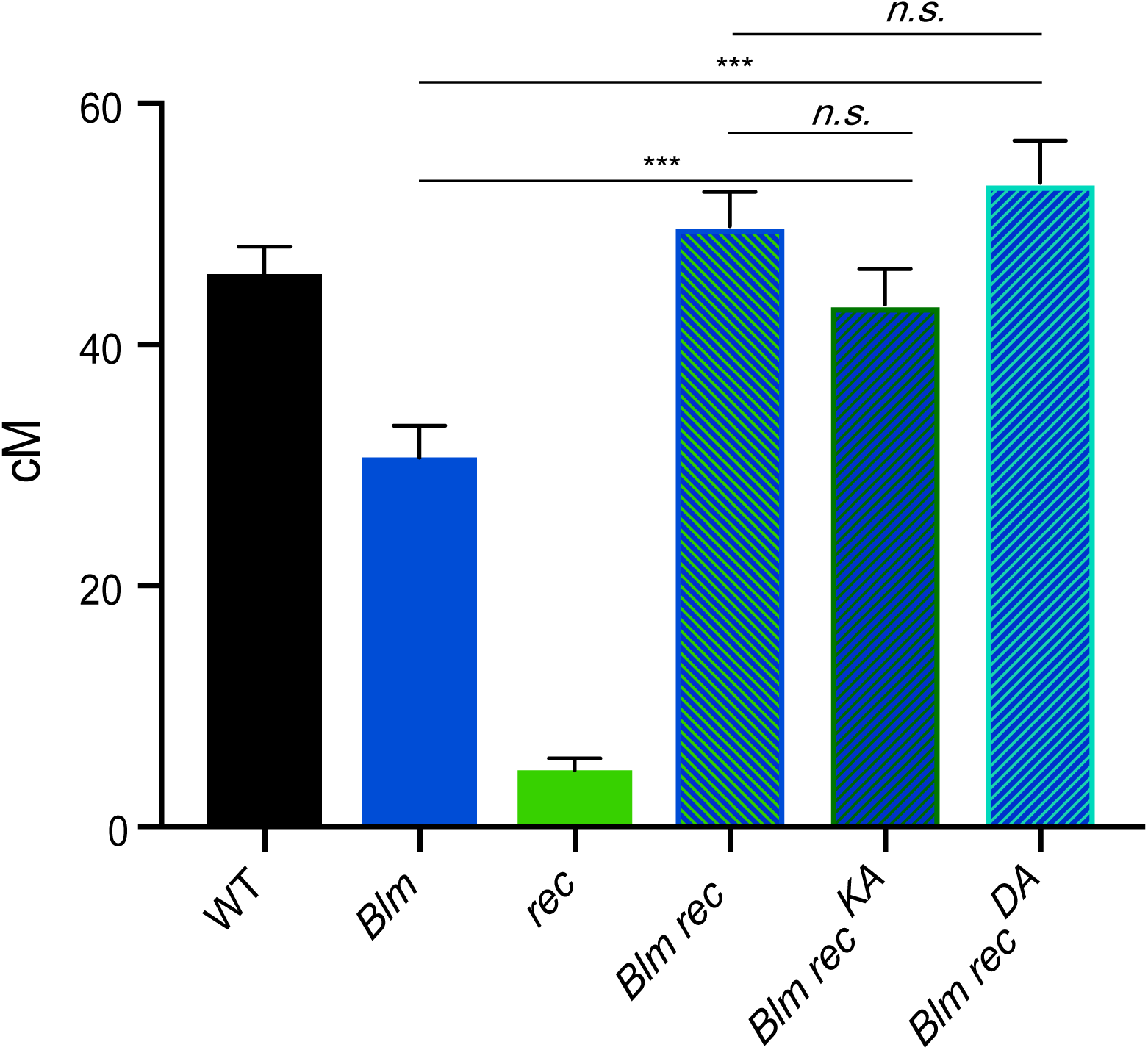
Requirements of REC ATPase activity in Blm function. Map units of *WT* (Hatkevich *et al.* 2017), *Blm* (Kohl *et al.* 2012), *rec, Blm rec* (Kohl *et al.* 2012), *Blm rec*^*KA*^, and *Blm rec*^*DA*^. Map units represented as centimorgans (cM). Error bars show 95% confidence intervals. Refer to Table S3 for complete data set. *** *p* < 0.0001. (*Blm rec* vs *Blm rec*^*KA*^ *p* = 0.10) (*Blm rec* vs *Blm rec*^*DA*^ *p* = 0.52).

In summary, the mei-MCMs are a family of diverged proteins that help to establish the recombination landscape in *Drosophila melanogaster* by promoting Class I crossovers and inhibiting Class II crossovers. Results obtained in this study have further elucidated meiotic recombination roles for two mei-MCM proteins, MEI-218 and REC. While the N-terminus of MEI-218 is dispensable for crossover formation (Figure 2), REC’s predicted ability to bind and hydrolyze ATP exhibit differential requirements for regulating Class I and Class II crossover formation. From our genetic analyses, we suggest that the Walker B motif of REC, but not the Walker A motif, is required for promoting the formation Class I, MEI-9 dependent crossovers (Figures 3 and 4). The weakly antimorphic phenotype of *rec*^*DA*^ demonstrates that an impaired REC Walker B mutant renders a poisonous complex – a complex in which we propose cannot be released from recombination sites. Both Walker A and Walker B motifs block crossovers in the Class II pathway, suggesting that REC forms different complexes to execute its pro- and anti-crossover functions. Biochemical and cytological studies are needed to support or refute these hypotheses.

## Acknowledgements

We thank Juan Carvajal Garcia, Carolyn Turcotte, and anonymous reviewers for thoughtful comments. This work was supported in part by a grant from the National Institute of General Medical Sciences to J.S. under award 1R35 GM-118127. K.P.K. was supported in part by NIH grant P20GM103499. T.H. was supported in part by NIH grants 5T32GM007092 and 1F31AG055157.

## Author Summary

Crossover formation between homologs is essential for accurate segregation at the end of meiosis I. Crossovers are typically formed through two pathways: Class I and Class II. The mei-MCMs are a class of proteins that promote Class I crossover formation and prohibit Class II crossovers in Drosophila. Although the mei-MCMs are conserved, little is known about their function. Here, we investigate the roles of two mei-MCMs, REC and MEI-218, in Class I and Class II crossover formation in Drosophila. From results in this study, we generate novel, testable hypotheses to further elucidate the meiotic function of the mei-MCM proteins.

