## Supplemental Figures for "Meiotic MCM proteins promote and inhibit crossovers during meiotic recombination"

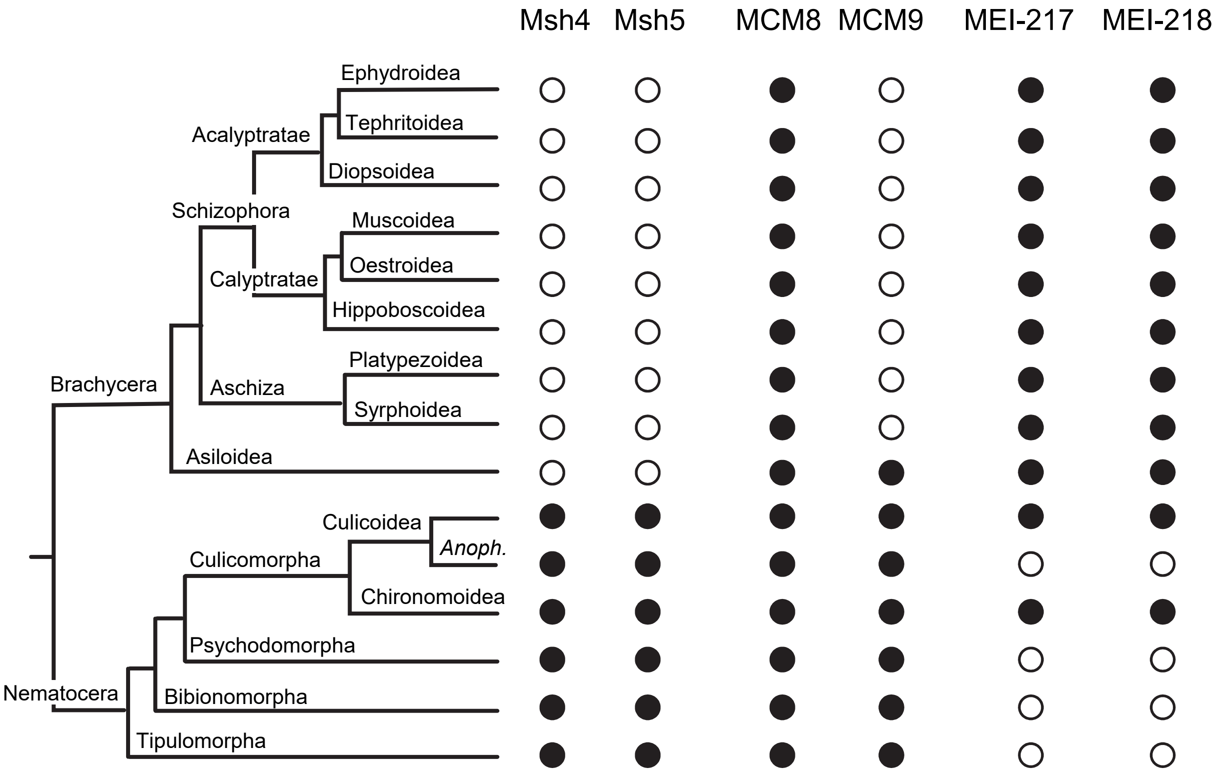


**Figure S1. Occurrence of Msh4, Msh5, MCM8, MCM9, MEI-217, and MEI-218 in Diptera.** The dendrogram on the left illustrates relationships among Dipteran taxa for which sufficient genome or transcriptome sequence is available to determine with reasonable confidence the presence or absence of genes encoding proteins relevant to this work. Circles to the right indicate presence (filled) or absence (open) of each gene/protein. For the suborder Brachycera, major superfamilies within Schizophora and the sister taxon Aschiza are shown, as well as the superfamily Asiloidea. For the suborder Nematocera, only infraorders are shown, except for Culicomorpha, where both superfamiles are indicated. Within the superfamily Culicoidea (mosquitoes), MEI-217 and MEI-218 are found in *Culex* and *Aedes* but are missing from all of the 20 *Anopheles* species whose genomes are sequenced.

It is hypothesized that the mei-MCM complex functionally replaces Msh 4/5 in *Drosophila* (Kohl, Jones, and Sekelsky 2012). We do not find orthologs of Msh4, Msh5, or Mcm9 in species in the Dipteran sub-order Brachycera, suggesting that the structure and function of the *Drosophila* mei-MCM complex may have its origins in the ancestral founder of this lineage. Interestingly, Asiloidea appear to have retained an ortholog of MCM9. It may be informative to examine these species more thoroughly when additional sequences become available.

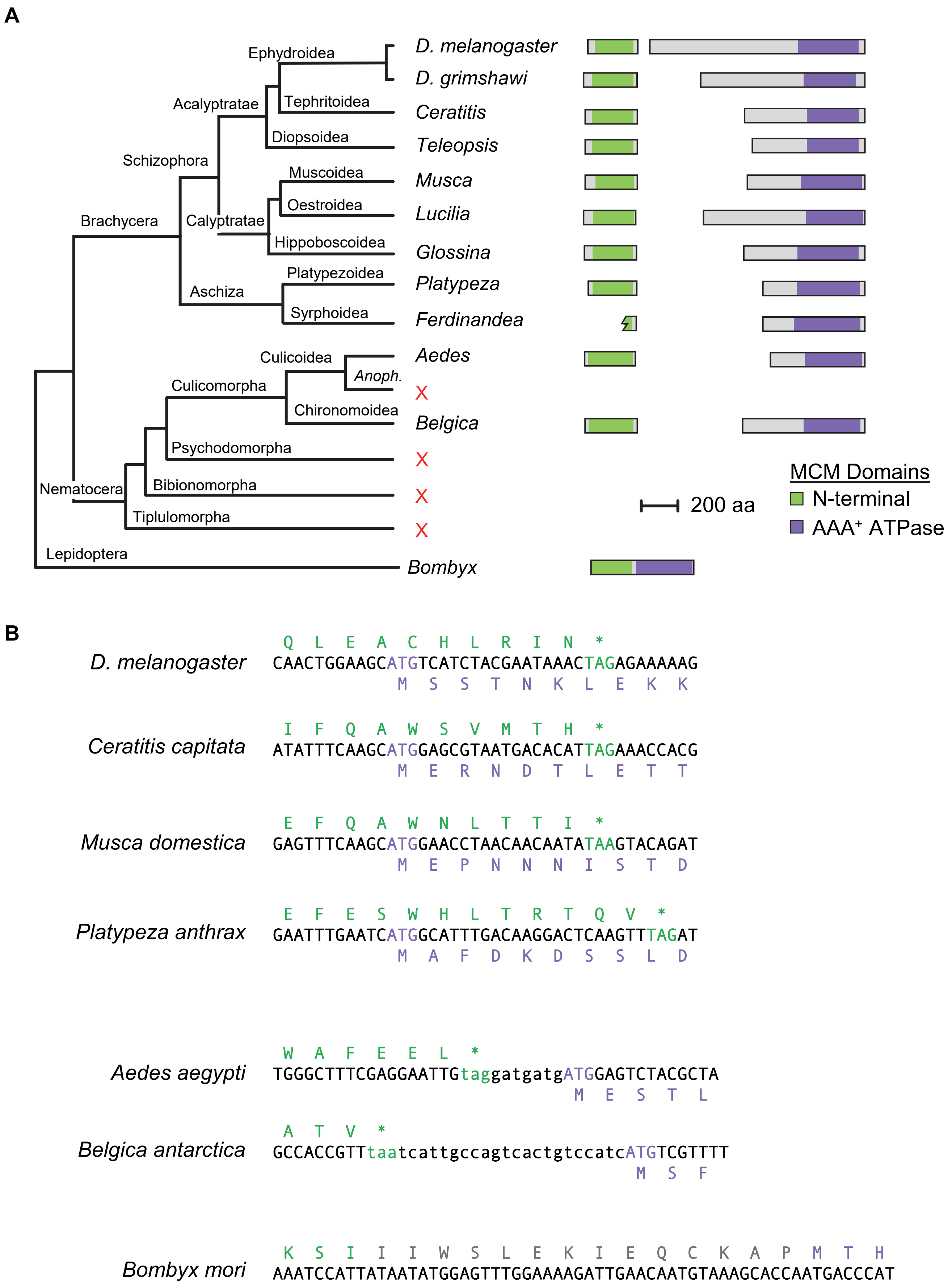
A
Ephydroidea
Tephritoidea
Acalyptratae
Diopsoidea
Schizophora
Muscoidea
Oestroidea
Brachycera
Calyptratae
Hippoboscoidea
Platypezoidea
Aschiza
Syrphoidea
Culicoidea
Anoph.
Culicomorpha
Chironomoidea
Psychodomorpha
Nematocera
Bibionomorpha
Tiplulomorpha
Lepidoptera
D. melanogaster
D. grimshawi
Ceratitis
Teleopsis
Musca
Lucilia
Glossina
Platypeza
Ferdinandea
Aedes
X
Belgica
X
X
X
Bombyx
200 aa
MCM Domains
N-terminal
AAA+ ATPase
B
D. melanogaster
Q L E A C H L R I N *
CAACTGGAAGCATGTCATCTACGAATAAACTAGAGAAAAAG
M S S T N K L E K K
Ceratitis capitata
I F Q A W S V M T H *
ATATTTCAAGCATGGAGCGTAATGACACATTAGAAACCACG
M E R N D T L E T T
Musca domestica
E F Q A W N L T T I *
GAGTTTCAAGCATGGAACCTAACAACAATATAAGTACAGAT
M E P N N N I S T D
Platypeza anthrax
E F E S W H L T R T Q V *
GAATTTGAATCATGGCATTTGACAAGGACTCAAGTTTAGAT
M A F D K D S S L D
Aedes aegypti
W A F E E L *
TGGGCTTTCGAGGAATTGtaggatgatgATGGAGTCTACGCTA
M E S T L
Belgica antarctica
A T V *
GCCACCGTTtaatcattgccagtcactgtccatcATGTCGTTTT
M S F
Bombyx mori
K S I I I W S L E K I E Q C K A P M T H
AAATCCATTATAATATGGAGTTTGGAAAAGATTGAACAATGTAAAGCACCAATGACCCAT

**C**

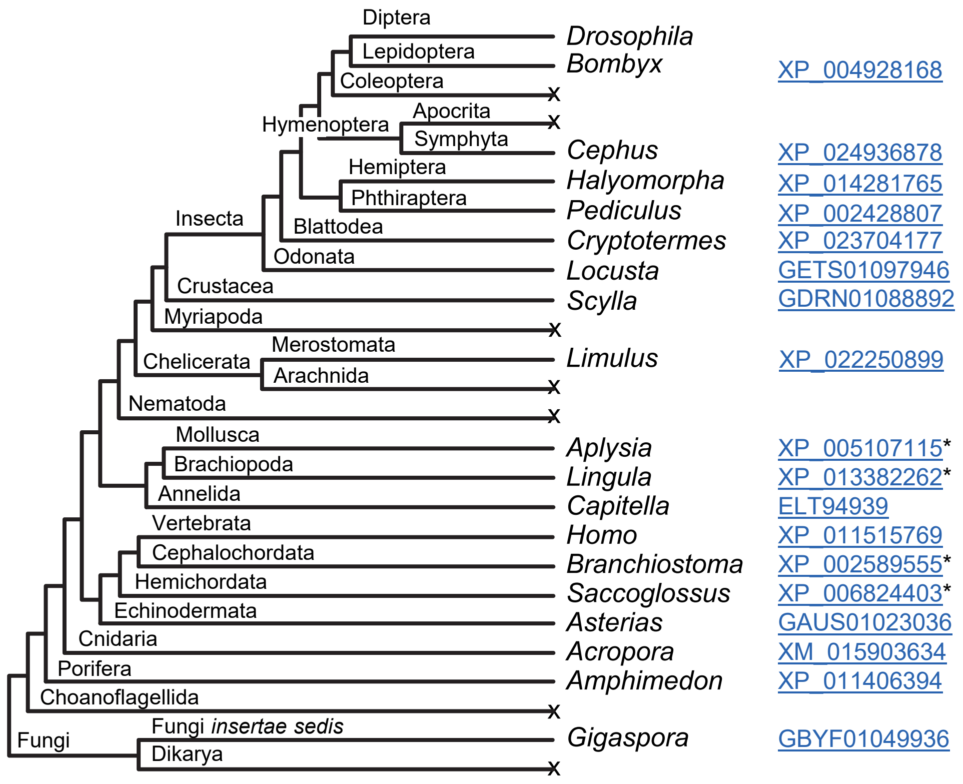
Diptera
Lepidoptera
Coleoptera
Hymenoptera
Apocrita
Symphyta
Hemiptera
Phthiraptera
Insecta
Blattodea
Odonata
Crustacea
Myriapoda
Merostomata
Chelicerata
Arachnida
Nematoda
Mollusca
Brachiopoda
Annelida
Vertebrata
Cephalochordata
Hemichordata
Echinodermata
Cnidaria
Porifera
Choanoflagellida
Fungi insertae sedis
Fungi
Dikarya
Drosophila
Bombyx
Cephus
Halyomorpha
Pediculus
Cryptotermes
Locusta
Scylla
Limulus
Aplysia
Lingula
Capitella
Homo
Branchiostoma
Saccoglossus
Asterias
Acropora
Amphimedon
Gigaspora
XP_004928168
XP_024936878
XP_014281765
XP_002428807
XP_023704177
GETS01097946
GDRN01088892
XP_022250899
XP_005107115*
XP_013382262*
ELT94939
XP_011515769
XP_002589555*
XP_006824403*
GAUS01023036
XM_015903634
XP_011406394
GBYF01049936

**Figure S2. Structures of MEI-217 and MEI-218 in Diptera.** (A) The dendrogram is the same as in Figure S1, with additional species to illustrate the variation in domain architectures. Domain architectures for representative species are to the right (the jagged N-terminal end in *Ferdiandea cuprea* indicates incomplete sequence). Domains were determined by PHYRE2 alignment to Protein Data Bank entry c5udb7 (a cryo-electron microscopy structure of *S. cerevisiae* MCM7). Accession numbers for the sequences included are listed below. Accession numbers that start with J are from Ensemble Metazoa genomic assemblies (found at http://metazoan.ensemble.org). The *Aedes*, *Musca*, and *Glossina* sequences are genomic contigs from Vectorbase (http://vectorbase.org). All other sequences are from NCBI (http://ncbi.nih.nlm.gov); those starting with a G are from the transcriptome shotgun assembly (TSA) database.

| Species | Accession |
|---|---|
| *Drosophila melanogaster* | NM_167557.3 |
| *Drosophila grimshawi* | XM_001992187.1 |
| *Ceratitis capitate* | GAMC01014250.1 |
| *Teleopsis whitei* | GBBQ01026862.1 |
| *Musca domestica* | scf7180000644883 |
| *Lucilia cuprina* | JRES01000755:1975-11359 |
| *Glossina morsitans* | scf7180000644883 |
| *Platypeza anthrax* | GCGU01008763.1 |
| *Ferdinandea cuprea* | GCHQ01011487.1 |
| *Aedes aegypti* | AAGE02016621.1 |
| *Belgica Antarctica* | JPYR01000187:32247-35225 |
| *Bombyx mori* | AK381112.1 |

(B) Junctions between open reading frames (ORFs) for the N-terminal and AAA+ ATPase domains are shown. At the top are three species from Shizophora and one from Aschiza, showing overlapping ORFs. Amino acids at the end of the N-terminal domain are shown in green above the DNA sequence (the position of the stop codon is highlighted in green); amino acids for the beginning of the AAA+ ATPase domain are in purple below the DNA sequence (the position of the start codon is highlighted in purple). Below that are two Nematocera species, showing separate but non-overlapping ORFs. Non-coding sequence between the ORFs is in lowercase text. At the bottom is a non-Diptera representative, the moth *Bombyx mori*. In this case, as in all other non-Dipteran species with an Mcmdc2 ortholog and in replicative MCM proteins, the two domains are on the same polypeptide, separated by a short linker.

(C) Distribution of Mcmdc2 in Opisthokonta. Dendrogram shows phyla, some sub-phyla, and several orders within the sub-phylum Insecta in which we can find clear orthologs of Mcmdc2 or in which there are sufficient genome or transcriptome sequences to suggest loss of Mcmdc2 with reasonable confidence. For those clades with an ortholog, a representative genus is listed, along with an accession number. Taxa in which we could not find any orthologs are indicated with an x. We have not found Mcmdc2 orthologs outside of Opisthokonta.

Figure S3 Hartmann *et al.*

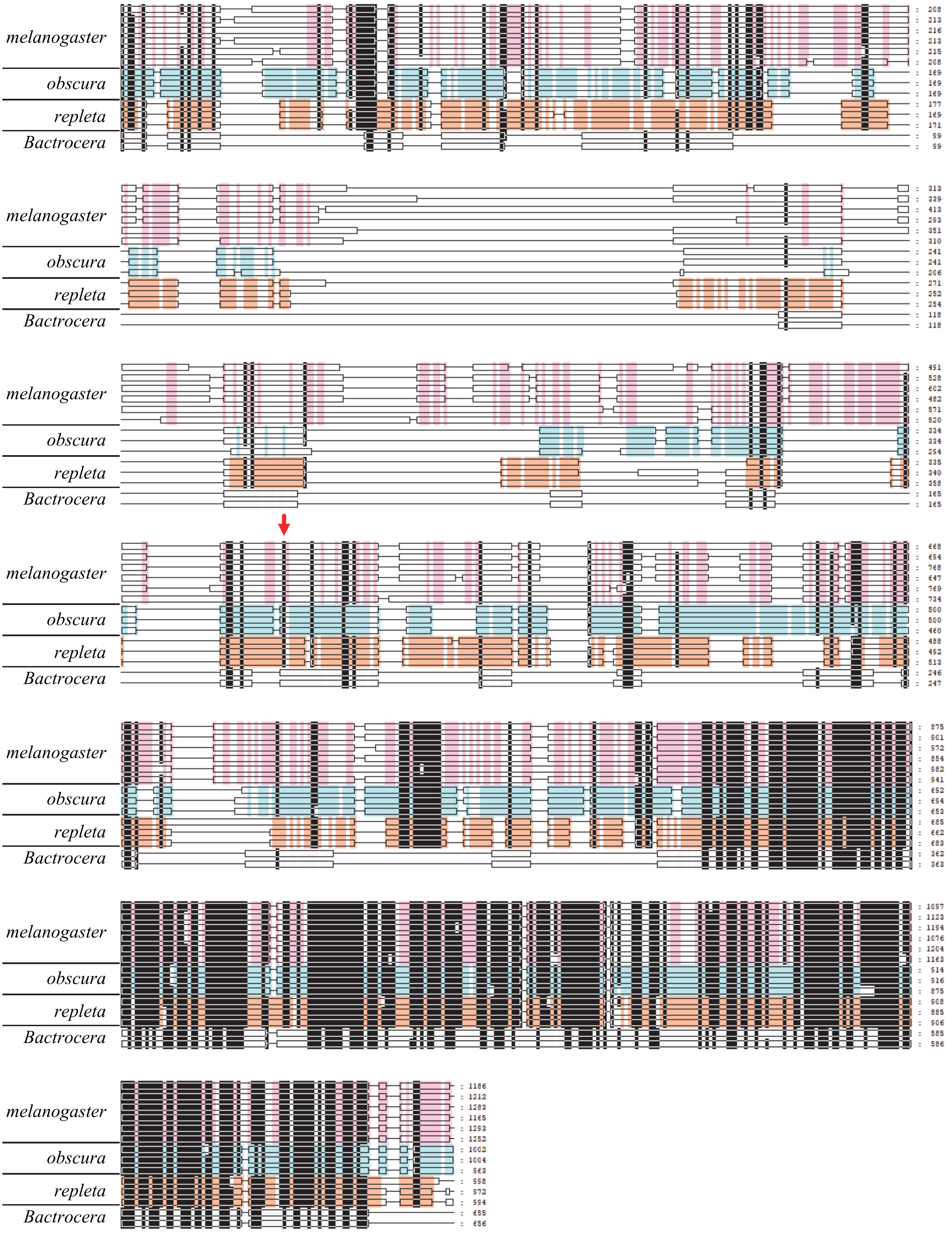

**Figure S3. Sequence conservation and divergence in MEI-218.** MEI-218 orthologs from 12 *Drosophila* species and two *Bactrocera* species (Tephritid fruit flies, also in the Acalyptratae subsection of Schizophora) are depicted. The red arrow indicates the start of the N-terminally-truncated MEI-218 analyzed in this work.

Species are, from top to bottom and sorted by species group as indicated on the figure:

| | |
|---|---|
| *melanogaster*: | *Drosophila melanogaster*<br>*Drosophila sechelia*<br>*Drosophila simulans*<br>*Drosophila mauritiana*<br>*Drosophila yakuba*<br>*Drosophila erecta* |
| *obscura*: | *Drosophila pseudoobscura pseudoobscura*<br>*Drosophila persimilis*<br>*Drosophila miranda* |
| *repleta*: | *Drosophila navojoa*<br>*Drosophila mojavensis*<br>*Drosophila arizonae* |
| *Bactrocera*: | *Bactrocera latifrons*<br>*Bactrocera dorsalis* |

Thin horizontal lines denote gaps introduced in the alignment process. Vertical lines indicate amino acid identity or similarity, using the Dayhoff PAM 200 matrix. Black is conserved among at least 11 sequences (*e.g.*, one mismatch in the 12 *Drosophila* species). Pink indicates conservation within the *melanogaster* species group, aqua within the *obscura* group, and orange within the *repleta* group. The red arrow denotes the start codon for the truncated MEI-218 described in the text.

Sequences were aligned in MEGA (v. 10.0.4) using the MUSCLE algorithm. Manual adjustments were done in GeneDoc v. 2.7.000. This visualization is the summary view produced by GeneDoc, with species groups with conservation mode shading enabled.

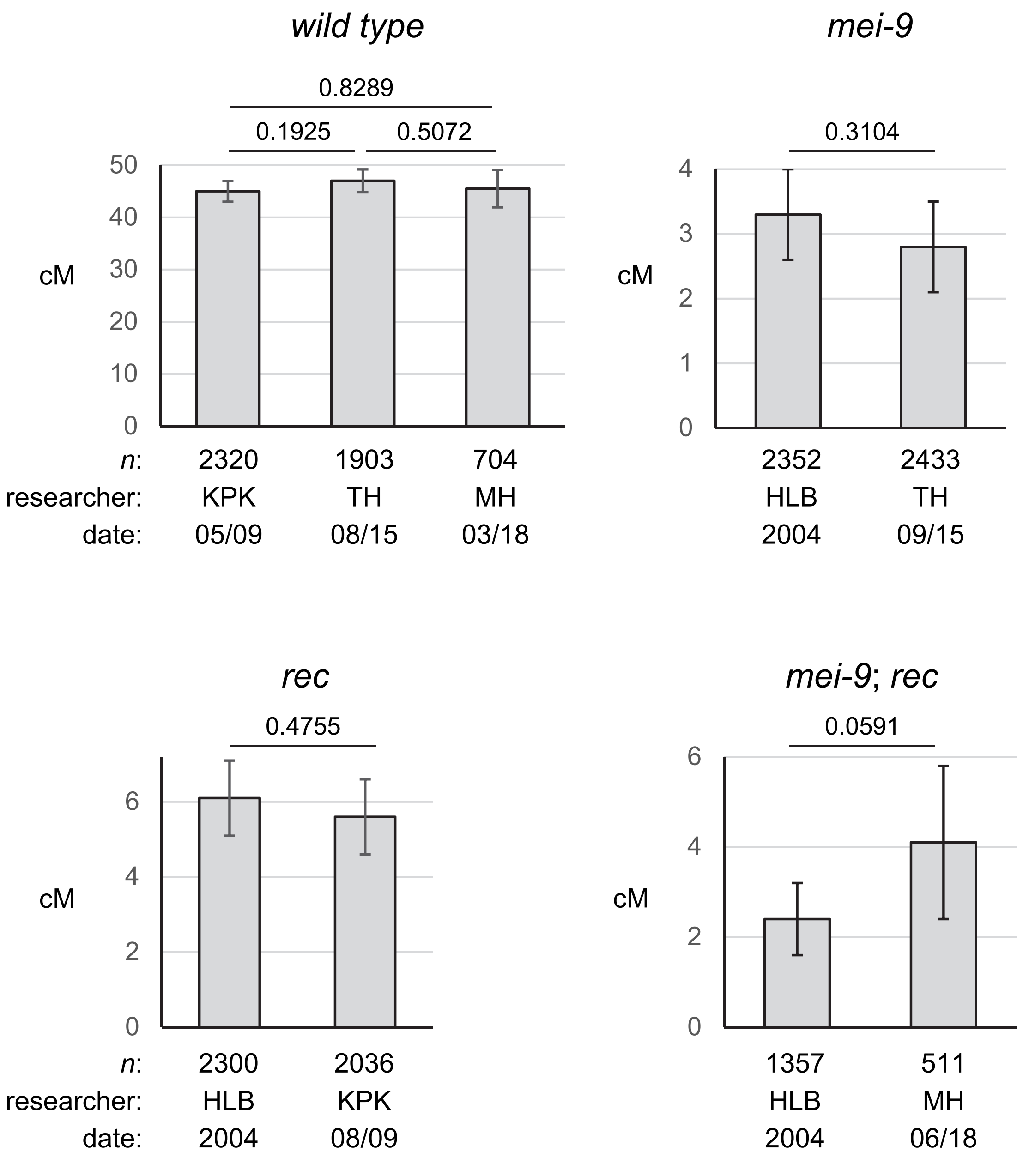


**Figure S4. Reproducibility of recombination assays.** Graphs shows the total number of cM for the *net* – *cn* intervals (*al* – *cn* for HLB assays; *al* is 0.2 cM to the right of *net*) for recombination assays in the same genotype scored by different researchers at different times. Error bars are 95% confidence intervals. *P* values are from Fisher's exact test on 2x2 contingency tables with total crossovers and parental flies for each genotype. Below each bar is the total number of flies scored (*n*), the researcher doing the scoring, and the date of the experiment. MH, KPK, and TH are authors of this work, with some data published in Kohl *et al.* 2012 and Hatkevich *et al*. 2017. HLB refers to data published in Blanton *et al.* 2005. The *mei-9*; *rec* MH experiment had six double crossovers and one triple crossovers. Because all of these were *b* $pr^{+}$ *cn* and no multiple crossover events were observed in the 2.7x larger HLB dataset, we suspect these were from some non-meiotic event(s); these flies were removed from this analysis.

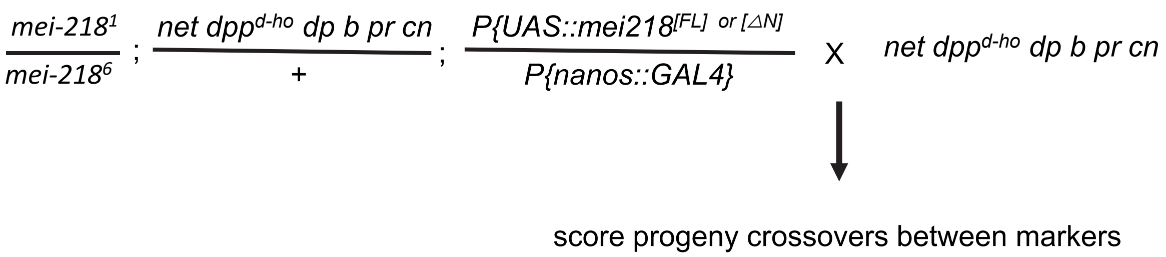


**Figure S5. Cross scheme of *mei-218* overexpression.** *mei-218*$^{FL}$ or *mei-218*$^{\Delta N}$ are expressed from *UAS::Gal4* system on the third chromosome in the background of a *mei-218* null mutation. Recessive markers on chromosome 2 are used for crossover scoring. Virgin female progeny are crossed back to flies homozygous for the recessive marker chromosome.
