## Supplemental Tables for "Meiotic MCM proteins promote and inhibit crossovers during meiotic recombination"

**Table S1. Meiotic crossovers on chromosome *2L* in *mei-218$^{FL}$* and *mei-218$^{\Delta N}$*.**

| Interval | *mei-218$^{FL}$* | *mei-218$^{\Delta N}$* | Fold Change | *P* Value |
|---|---|---|---|---|
| *net-ho* | 4.87 (1.53) | 5.9 (1.08) | 1.21 | n.s. |
| *ho-dp* | 5.79 (1.66) | 7.49 (1.21) | 1.29 | <0.0001 |
| *dp-b* | 25.39 (3.1) | 21.87 (1.91) | 0.86 | <0.0001 |
| *b-pr* | 11.71 (2.29) | 13.55 (1.58) | 1.16 | n.s. |
| *pr-cn* | 6.32 (1.73) | 7.11 (1.18) | 1.13 | n.s. |
| Total *n* | 759 | 1815 | | |

Crossover frequency shown as map units (centimorgans). Numbers in parentheses represent 95% confidence intervals. Fold change = *mei-218$^{FL}$* / *mei-218$^{\Delta N}$*. *P* values determined by chi-squared tests. Figure 2 shows distributions for *mei-218$^{FL}$* and *mei-218$^{\Delta N}$*.

**Table S2.Nondisjunction of *WT*, *rec$^{-}$* / *rec$^{+}$*, and *rec$^{DA}$* / *rec$^{+}$*.**

| Genotype | *XX* females | *XY* males | *XXY* females | *XO* males | Total *n* | NDJ |
|---|---|---|---|---|---|---|
| *wild type (y w)* | 1551 | 1481 | 0 | 1 | **3034** | 0.07% |
| *rec$^{-}$* / *rec$^{+}$* | 880 | 775 | 0 | 0 | **1655** | 0% |
| *rec$^{DA}$* / *rec$^{+}$* | 1072 | 863 | 10 | 4 | **1963** | 1.43% |

*XX* females and *XY* males are normal, whereas *XXY* females and *XO* males are karyotypes genotypes resulting from nondisjunction. Total *n* calculates in exceptional progeny that do not survive (*XXX* and *XO*).

**Supplemental Table S3. Crossovers in each interval on chromosome *2L* for all mutants discussed.**

| | Progeny | Genotype Being Assayed | | | | | | | | | | | | | | | |
|---|---|---|---|---|---|---|---|---|---|---|---|---|---|---|---|---|---|
| | | *WT* | *mei-218* | *mei-218* $^{FL}$ | *mei-218* $^{\Delta N}$ | *rec* | *rec*$^{KA}$ | *rec*$^{DA}$ | *rec*$^{DA}$ / *rec*$^{+}$ | *mei-9* | *mei-9; rec* | *mei-9; rec*$^{KA}$ | *mei-9; rec*$^{DA}$ | *Blm* | *Blm rec* | *Blm rec*$^{KA}$ | *Blm rec*$^{DA}$ |
| | Parental | 2376 | 1693 | 396 | 942 | 2129 | 1061 | 3452 | 1597 | 2366 | 490 | 3650 | 1360 | 844 | 705 | 684 | 445 |
| SCO | 1 (*net-ho*) | 176 | 4 | 24 | 75 | 8 | 54 | 3 | 73 | 6 | 0 | 7 | 0 | 23 | 31 | 27 | 13 |
| | 2 (*ho-dp*) | 290 | 6 | 33 | 93 | 10 | 100 | 5 | 137 | 11 | 3 | 13 | 1 | 22 | 29 | 35 | 10 |
| | 3 (*dp-b*) | 1099 | 16 | 168 | 315 | 53 | 464 | 29 | 679 | 40 | 11 | 43 | 5 | 89 | 136 | 116 | 98 |
| | 4 (*b-pr*) | 154 | 18 | 65 | 168 | 13 | 94 | 12 | 201 | 8 | 5 | 20 | 5 | 104 | 103 | 95 | 78 |
| | 5 (*pr-cn*) | 39 | 7 | 29 | 80 | 20 | 44 | 6 | 30 | 2 | 2 | 19 | 4 | 61 | 87 | 78 | 64 |
| DCO | 1 and 2 | 1 | 0 | 0 | 0 | 0 | 0 | 0 | 0 | 0 | 0 | 0 | 0 | 1 | 0 | 4 | 0 |
| | 1 and 3 | 11 | 0 | 3 | 13 | 0 | 7 | 0 | 4 | 0 | 0 | 0 | 0 | 4 | 11 | 4 | 0 |
| | 1 and 4 | 10 | 0 | 4 | 11 | 0 | 4 | 0 | 2 | 0 | 0 | 0 | 0 | 2 | 7 | 0 | 3 |
| | 1 and 5 | 2 | 0 | 6 | 8 | 0 | 0 | 0 | 2 | 0 | 0 | 0 | 0 | 1 | 0 | 3 | 2 |
| | 2 and 3 | 6 | 0 | 3 | 13 | 0 | 3 | 0 | 3 | 0 | 0 | 0 | 0 | 1 | 5 | 1 | 6 |
| | 2 and 4 | 7 | 0 | 6 | 18 | 0 | 1 | 0 | 9 | 0 | 0 | 0 | 0 | 2 | 7 | 5 | 2 |
| | 2 and 5 | 13 | 0 | 1 | 12 | 0 | 2 | 0 | 3 | 0 | 0 | 0 | 0 | 2 | 2 | 1 | 4 |
| | 3 and 4 | 19 | 0 | 10 | 38 | 0 | 14 | 0 | 14 | 0 | 0 | 0 | 0 | 5 | 19 | 5 | 18 |
| | 3 and 5 | 17 | 0 | 8 | 18 | 0 | 4 | 0 | 7 | 0 | 0 | 0 | 0 | 7 | 21 | 17 | 12 |
| | 4 and 5 | 2 | 0 | 2 | 11 | 0 | 3 | 0 | 3 | 0 | 0 | 0 | 0 | 2 | 8 | 9 | 10 |
| | TCO | 0 | 0 | 1 | 0 | 0 | 0 | 0 | 0 | 0 | 0 | 0 | 0 | 1 | 10 | 7 | 7 |
| | Total *n* | **4222** | **1744** | **759** | **1815** | **2233** | **1855** | **3507** | **2764** | **2433** | **511** | **3752** | **1375** | **1171** | **1181** | **1091** | **774** |

Each row shows the number of parental (P), single (SCO), double (DCO), and triple (TCO) crossovers for each genotype and each mutant discussed in the article. Total *n* represents all parental and recombinant flies scored for each genotype. Wild-type data are from Hatkevich *et al.* 2017. Data for *mei-218 Blm*, and *Blm rec* are from Kohl, Jones, and Sekelsky 2012. The *mei-9*; *rec* experiment had six apparent double crossovers and one triple crossovers. Because all of these were *b pr*$^{+}$ *cn* and no multiple crossover events were observed in the 2.7x larger dataset of Blanton *et al.* (2005), we suspect these were from some non-meiotic event(s); these flies were therefore removed from this analysis.
